# Levô in fimo test – A simple test for fecal microbial reconstituted mice

**DOI:** 10.1101/2020.05.11.089581

**Authors:** SM Musheer Aalam, Scott I Gamb, Purna Kashyap, Nagarajan Kannan

**Affiliations:** Division of Experimental Pathology, Department of Laboratory Medicine and Pathology, Mayo Clinic, Rochester, Minnesota, 55905, USA; Microscopy and Cell Analysis Core, Mayo Clinic, Rochester, Minnesota, 55905, USA; Department of Gastroenterology, Mayo Clinic, Rochester, Minnesota, 55905, USA; Center for Regenerative Medicine, Mayo Clinic, Rochester, Minnesota, 55905, USA; Mayo Clinic Cancer Center, Mayo Clinic, Rochester, Minnesota, 55905, USA

## Abstract

Here we report the first method for rapid on-site monitoring of germfree C57BL/6 mice colonized with feces from conventionally raised mice. The method, named *levô in fimo* test, readily distinguishes feces from germfree mice and conventionalized germfree mice based on its properties to float or sink in Trump’s fixative solution. We validated the test using feces from conventionally raised mice, germfree mice and conventionalized germfree mice, and cross-validated using multiple indicators of microbial presence in feces. Levô in fimo test is a simple, cost-effective, instant yet robust method for on-site monitoring of gut microbial reconstitution in germfree mice.

## Main

The naturally excreted feces is a transient habitat composed of a complex mixture of non-living and living systems, where living systems maintain a homeostatic relationship with each other, share non-living resources, and relate symbiotically with their host^1^. The bulk of living systems in feces is constituted by bacteria, archaea and fungi^2^. The feces of ‘germfree’ mice lack detectable units of microorganisms^3^. The experimental procedure to transplant donor feces to reconstitute the recipient animal with a complex microbial constituents are now routinely used in many laboratories to understand host-microbial relationship to gain insight into various health benefits and diseases^4^. Research involving germfree mice is labor-intense and expensive. Herein we describe lev<u>ô</u> in fimo test, a new simple, cost-effective and foolproof method for on-site monitoring of germfree mice during peri-fecal transplantation window.

We have compared feces collected from three groups of C57BL/6 mice (Taconic Biosciences) and they are (a) conventionally raised mice (henceforth referred to as germ mice), (b) mice raised in germfree isolators (germfree mice), or (c) germfree mice treated with a single oral gavage of 300 μl preparation of previously frozen feces from donor germ mice (see Methods). The germfree mice 3-weeks post-fecal transplantation is henceforth referred to as ‘feces transplanted-germfree mice’.

For lev<u>ô</u>in fimo test, feces from the three groups of mice, were dropped into Trump’s fixative solution^5^ in a clear tube and gently agitated to submerge the feces into the fixative solution. The observations, whether the feces floats or sinks to the bottom of the solution, were instantly (< 1 minute) recorded. The results were as follows. All feces from germfree mice sank instantly (Figure 1A). Inversely, all feces from germ mice and feces transplanted-germfree mice floated in Trump’s fixative solution (Figure 1A). These findings strongly indicated that the fecal floatation behavior in Trump’s fixative solution was directly associated with complex microbial constituents in the feces. To further validate the association between fecal behavior in Trump’s fixative and fecal microbial colonization, we used multiple methods that indicate microbial presence in fecal samples. First, we examined fecal samples from all three cohorts of mice using scanning electron microscopy (SEM). At higher resolution diverse microorganisms were evident in feces from germ mice and feces transplanted-germfree mice whereas germfree mice had no evidence of microorganisms (Figure 1B). Interestingly, microorganisms were more uniformly distributed in germ mice compared to feces transplanted-germfree mice (Figure 1B). Secondly, we isolated DNA for bacterial PCR from feces from all three groups. Total fecal DNA measurements in feces showed that germ and feces transplanted-germfree mice had similar levels i.e., ∼32μg and ∼30μg respectively per mg wet weight of feces (Figure 1C). Total fecal DNA in germfree mice was around ∼2-3μg per mg wet weight of feces, a ∼15-fold reduction compared to other two groups (Figure 1C). Thirdly, we used PCR methods to detect microbial presence^6, 7^. PCR confirmed presence of bacterial presence in germ mice and feces transplanted-germfree mice and absence in germfree mice (Figure 1D). Additionally, feces of germfree mice were placed in Sabouraud dextrose media, brain-heart infusion media, and nutrient broth media at 37°C for 7 days under aerobic and anaerobic conditions. No growth activity of microorganism was a further confirmation of germfree status of feces in our mice (data not shown). Therefore, the floatation or sinking properties of feces in Trump’s fixative solution detected by levô in fimo test (Figure 1A) was directly linked to the presence or absence complex microbial living systems respectively. Henceforth, it would be indicated that levô in fimo test’s positive for complex microbial presence if the feces instantly floats in Trump’s fixative, and negative if it instantly sinks.

**Figure 1:**
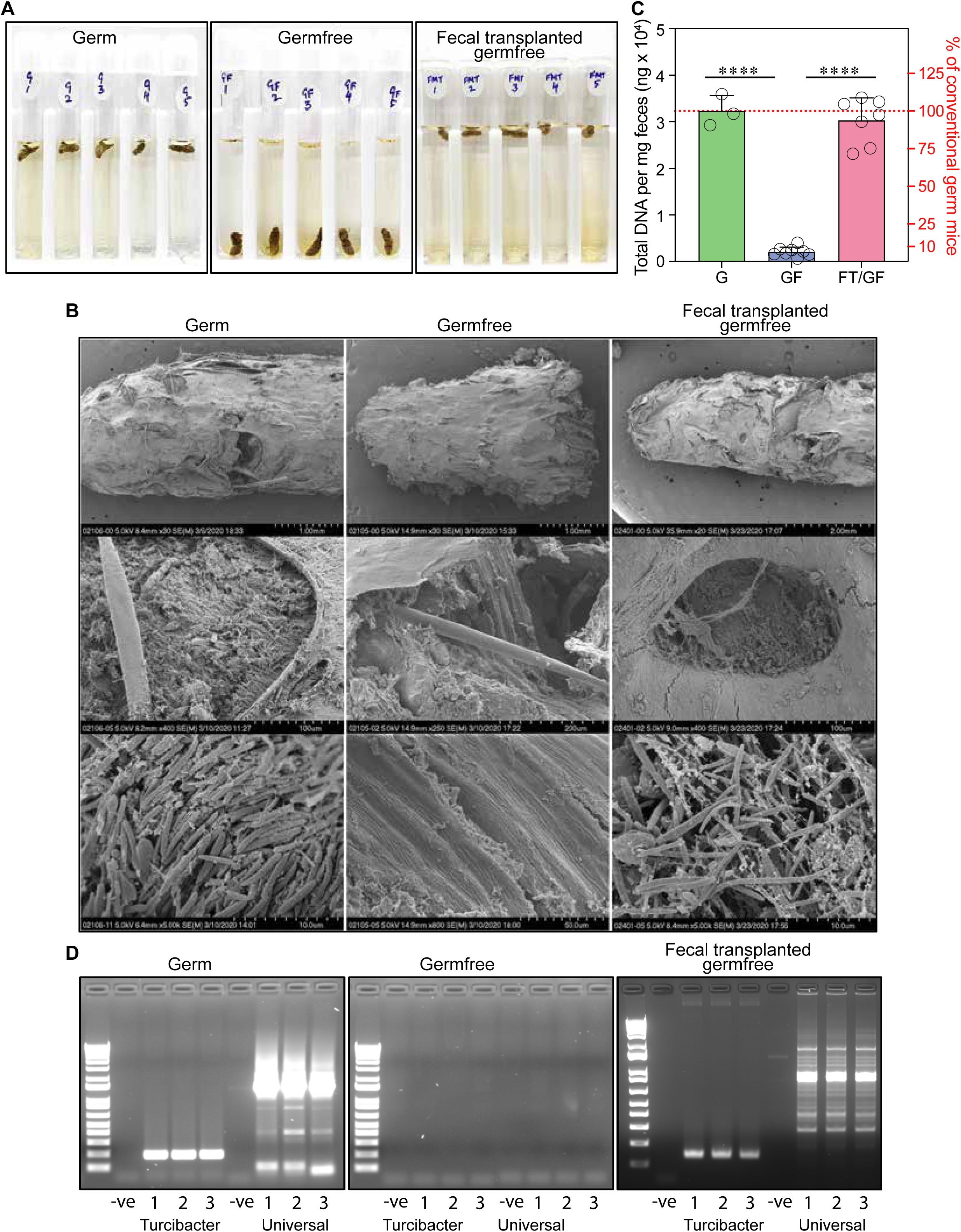
Levô in fimo test and validation. A) Levô in fimo tests in feces obtained from germ, germfree and feces transplanted-germfree mice (3 week post-transplant). B) Representative SEM images of feces at three different resolutions. C) Total fecal DNA levels per mg wet weight of feces. **** represents Fisher’s exact test p < 0.0001. D) Bacterial PCR test results.

We then tested if levô in fimo test could be used for on-site monitoring of gut microbial reconstitution kinetics in germfree mice during peri-fecal transplantation window. In a proof-of-concept experiment, we collected feces from germfree mice 0-2 hour before fecal transplantation and then at 1-3 weeks post-fecal transplantation. Each fecal sample was split into two and used for levô in fimo test and total fecal DNA measurement (an indicator of microbial load) (Figure 2A). All fecal samples in pre-feces transplanted-germfree mice tested negative (Figure2B) and the results correlated with fecal DNA levels (Figure 2C). Test was100% positive for fecal samples at all timepoints post-fecal transplant. The corresponding DNA levels suggested that following fecal transplantation, microbes multiplied extensively, reached a steady-state with host and persisted at high levels (Figure 2C). Fecal DNA levels during pre-fecal transplantation timepoint in germfree mice were consistently around ∼2-3μg per mg wet weight of feces and all post-fecal transplantation timepoints had ∼10-15 fold higher DNA levels reaching DNA levels observed in germ mice (Figure 1C) indicating that 100% of germfree mice were conventionalized i.e., microbially colonized by feces from conventionally raised mice.

**Figure 2:**
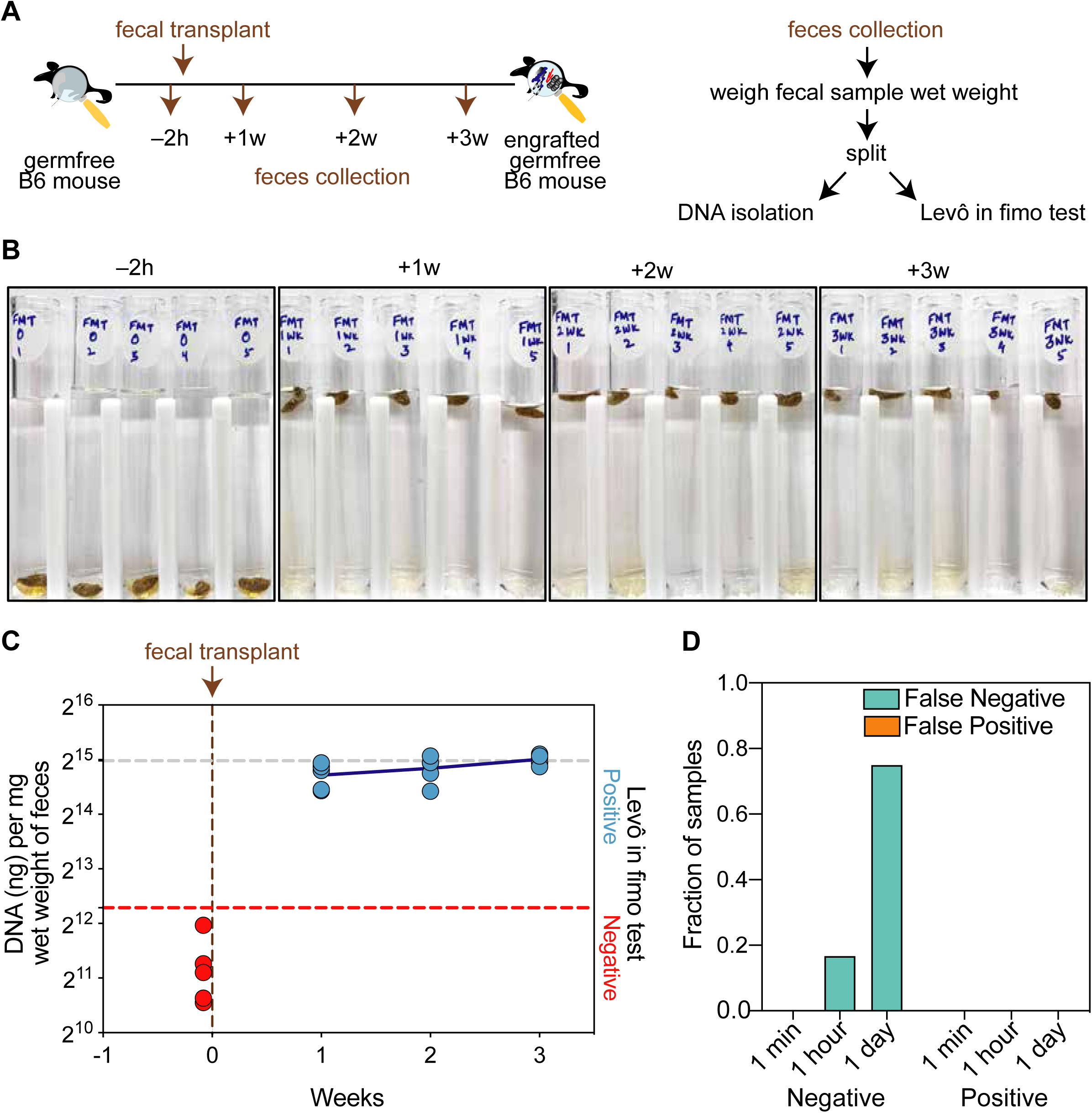
Levô in fimo test and gut microbial reconstitution kinetics. A) Schematics. B) Images of levô in fimo test results at 1 min. D) Fecal reconstitution kinetics in germfree mice during peri-fecal transplantation window. E) Levô in fimo test efficacy over time.

To determine the efficacy of levô in fimo test we let the feces from above experiment (Figure 1C) remain in solution for extended durations i.e., 1 hour and 1 day and recorded test results (Figure 1D). While 1 minute results showed 0% false positive or false negative test, were observed time-dependent increase in false negative results. Therefore, it is essential that levô in fimo test using fecal samples need to be recorded instantly (in seconds to minutes and not hours) to ensure 100% accuracy.

Levô in fimo test can be used for on-site at point-of-animal care using feces freshly excreted or from animal cage and off-site using frozen feces to monitor gut microbial reconstitution in germfree mice. The other advantages include the test is most simple to learn, per test cost is extremely economical (<$1/test) and does not require machines. The test results are instant (<1 minute) which save research time, effort and operational cost. Future applications of levô in fimo test on feces collected from different strains of germfree mice on different dietary and with pathological conditions following complex (fecal) or simple (gnotobiotic) gut microbial reconstitutions from different donors could be explored.

## Methods

### Germ and germfree mice

Female 6-8 week old conventional (referred as germ) C57BL/6NTac (B6) mice were obtained from Taconic Biosciences and maintained in standard mouse facility. Mayo Clinic Germfree Facility bred and maintained germfree B6 mice (originally obtained from Taconic Biosciences) in sterile isolators (Class Biologically Clean, Ltd.). Age-matched female germfree B6 mice were used in this study. Germ mice were fed with Picolab® mouse diet 20 5058 Germfree mice were fed an autoclaved standard diet (LabDiet® 5K67). All procedures were reviewed and approved by Mayo Clinic Institutional Animal Care and Use Committee.

### Fecal sample collection

Freshly excreted feces were collected from the cages of germ, germfree and feces transplanted germfree mice once a week during cage changes. The fecal samples were either immediately used or frozen at –80°C.

### Fecal transplantation in germfree mice

A fecal pellet of freshly collected and frozen from cages of germfree B6 mice was used to gavage 2 mice. In brief, a fecal pellet was homogenized in 750μl pre-reduced PBS in anaerobic COY chamber and 300 μl of fecal suspension was loaded into 1 ml syringes attached with gavage needles. Subsequently, the mice were restrained by hand and the gavage needle was inserted through the esophagus into the stomach, and fecal suspension was delivered by depressing the syringe plunger. The needle was removed and the mouse was placed back into the cage. All the mice received fecal preparation from single donor cage.

### Monitoring of gnotobiotic isolator and mice using anaerobic and aerobic microbial culture method

Fecal pellets were placed into a clean culture hood and added to two sets of tubes with either calf brain-beef heart infusion broth medium for the growth of aerobic bacteria, nutrient broth medium fastidious and non-fastidious microorganisms or Sabouraud dextrose broth medium for the cultivation of yeast and molds ^1^. One set of tubes with each broth was placed into the incubator inside of the COY anaerobic chamber. The other set with each broth was placed into an std aerobic incubator. Samples were checked daily for a week for microbial growth. Gnotobiotic isolator and mice were referred to as ‘germfree’ if all tubes tested by this method remained clear during this period.

### Levô in fimo test

Freshly, collected fecal samples from germ and germfree B6 cages were dropped into 4 ml of Trump’s fixative solution in 5 ml clear tubes. The samples were observed for floatation at 1 min, 1 hour and 1 day later and images were captured.

### Scanning Electron Microscopy (SEM)

The fecal samples were fixed at 4°C for a day in Trumps fixative, 4% formaldehyde + 1% glutaraldehyde in a phosphate buffer ^2^. Samples were then washed in phosphate buffer, rinsed in water, and dehydrated through a graded series of ethanols (10%-30%-50%-70%-905-95%-100%×2). Then critical point dried (EMS) using carbon dioxide, mounted on an aluminum stub using conductive carbon SEM tabs (EMS), and sputter-coated with gold-palladium for 150 seconds. Finally, they were imaged in a Hitachi S-4700 cold field emission scanning electron microscope SEM (paired upper and lower secondary electron (SE) detectors) at 5kV accelerating voltage. All micrographs were acquired as TIFF images.

### Fecal DNA isolation

Fecal DNA was extracted using Masterpure™ complete DNA and RNA purification kit (MC85200) following manufacturer’s instructions with some modifications. In brief, 10 mg of fecal pellet was transferred to a tube with 300 μl tissue and lysis solution containing proteinase K and homogenized using Biomasher Ii® Closed System Disposable Micro Tissue Homogenizer. Samples were then incubated at 65ºC for 1 hour with agitation in Eppendrof Thermomixer C. The samples were cooled, centrifuged and the supernatant was transferred into a fresh tube and treated with RNase A at 37ºC for 30 mins. Subsequently, MPC protein precipitation solution was added to the samples, centrifuged and DNA was precipitated from the supernatant using isopropanol. The precipitated DNA was washed twice with 70% ethanol and resuspended in TE buffer. The yield of extracted DNA was estimated using Implen NanoPhotometer NP80. For engraftment kinetics, fecal DNA was extracted at different time points (2 hr prior to fecal transplant and then 1, 2 and 3 weeks post-fecal transplant) from excreted feces obtained during cage changes.

### Microbiome analysis by PCR

Microbiome analysis was performed with 500 ng of fecal DNA using 5 PRIME Hot Master Mix and Universal (Forward 5’-AGA GTT TGA TCC TGG CTC AG and reverse 5’-GAC GGG CGG TGW GTR CA) and Turicibacter (Forward 5’-GCG CGC AGG TGG TTA ATT AAG TCT and reverse 5’-TCA GTG TCA GTT GCA GAC CAG GAA) primers as described elsewhere^3,4^.

